# The effector recognition by synthetic sensor NLR receptors requires the concerted action of multiple interfaces within and outside the integrated domain

**DOI:** 10.1101/2022.08.17.504349

**Authors:** Xin Zhang, Yang Liu, Guixin Yuan, Dongli Wang, Tongtong Zhu, Xuefeng Wu, Mengqi Ma, Liwei Guo, Hailong Guo, Vijai Bhadauria, Junfeng Liu, You-Liang Peng

## Abstract

Plant sensor nucleotide-binding leucine-rich repeat (NLR) receptors detect pathogen effectors through their integrated domains (IDs). The RGA5 sensor NLR recognizes its corresponding effectors AVR-Pia and AVR1-CO39 from the blast fungus *Magnaporthe oryzae* through direct binding to its heavy metal-associated (HMA) ID to trigger the RGA4 helper NLR-dependent resistance in rice. Here we report a mutant of RGA5 named RGA5^HMA5^ that confers complete resistance in transgenic rice plants to the *M. oryzae* strains expressing the noncorresponding effector AVR-PikD. RGA5^HMA5^ carries three engineered interfaces, two of which lie in the HMA ID and the other in the C-terminal Lys-rich stretch tailing the ID. However, the RGA5 variants having one or two of the three interfaces, including replacing all the Lys residues with Glu residues in the Lys-rich stretch, failed to activate RGA4-dependent cell death of rice protoplasts. Altogether, this work demonstrates that sensor NLRs require a concerted action of multiple surfaces within and outside the IDs to both recognize noncorresponding effectors and activate helper NLR-mediated resistance, and has implications in structure-guided designing of sensor NLRs.

## Introduction

Many gene-for-gene diseases, including rice blast caused by *Magnaporthe oryzae*, pose a serious threat to global crop production and food security^1^. To cause such diseases, pathogens secrete a diverse array of effectors into host cells to subdue plant immunity^2^. To counter the effectors, plants have evolved nucleotide-binding leucine-rich repeat (NLR) immune receptors that recognize the avirulence (Avr) effectors either directly through physical binding or indirectly by monitoring the effector-mediated modification of guardee or decoy proteins and activate downstream immune responses^1^. Some of these NLRs function as sensor receptors that are genetically linked and physically paired with helper NLRs. Both sensor and helper NLRs share a tripartite domain architecture: an N-terminal coiled-coil (CC) or Toll/Interleukin −1 receptor domain^3^, a central nucleotide-binding (NB-ARC) domain and a C-terminal leucine-rich repeat (LRR) domain^4–6^. However, sensor NLRs usually carry an additional non-canonical integrated domain (ID; e.g., heavy metal-associated [HMA] or WRKY domains) that serves as “bait” to entice pathogen effectors^7–11^. Therefore, these IDs act as excellent targets for molecular engineering to create novel sensor NLR receptors.

Several studies have recently reported synthetic sensor NLRs with extended or altered effector recognition specificities via molecular engineering of the HMA IDs of Pik1 and RGA5. RGA5/RGA4 and Pik-1/Pik-2 are two paired NLR receptors in rice to confer blast resistance, within which RGA5 and Pik-1 function as the sensor NLRs, and RGA4 and Pik-2 as the helper NLRs. Notably, both RGA5 and Pik-1 carry an HMA ID for recognizing their corresponding ***M**agnaporthe* **A**vrs and To**x**B-like (MAX) effectors from *M. oryzae* ^8, 10, 12^. However, the two NLR systems differ in their working mechanisms. Pik-1 binds to its corresponding MAX effector AVR-PikD via an interface comprising the β2-β3-β4 sheet in the HMA ID while RGA5 physically interacts with the MAX effectors AVR-Pia and AVR1-CO39 through an interface composed of the α1 helix and β2 strand in the HMA ID^8, 10, 13, 14^. Rice germplasm contains multiple alleles of *Pik1*, which possess distinct effector recognition specificities and are polymorphic, mainly in their HMA IDs. For instance, Pikp-1 only recognizes AVR-PikD, while Pikm-1 can perceive AVR–PikD and two additional AVR-Pik variants (AVR-PikA and AVR-PikE)^15^. An engineered Pikp1 carrying the AVR-Pik binding interface of Pikm-1-HMA gained the capacity of Pikm1 to recognize the AVR-Pik variants, phenomimicking Pikm-1-mediated cell death in *N. bethamiana*^16^. By engineering the AVR1-CO39 interface of RGA5, we recently generated a designer NLR receptor RGA5^HMA2^ that confers specific resistance in transgenic rice to the *M. oryza*e strains expressing AVR-Pib^17^. However, Cesari *et al*. recently reported that the RGA5 mutants carrying the AVR-PikD binding interface of Pikp1-HMA can interact with the noncorresponding effector AVR-PikD but are unable to confer rice resistance to the *M. oryza*e strains expressing AVR-PikD^18^. Further, Wang et al. generated an RRS1 variant by introducing the *Phytoplasma* effector SAP05-dependent degron domain to the C-terminus of RRS1. This synthetic RRS1 receptor can recognize SAP05 but is uable to confer full resistance in transgenic Arabidopsis against the *Phytoplasma*^19^. These studies raise a key question as to why *de novo* effector-binding results in different outcomes in NLR activation in the host plants. We reason that *de novo* effector binding per se is necessary but insufficient for designer sensor NLRs to trigger immune responses in their host plants. Therefore, this study set out to define interfaces in the RGA5 HMA ID and its adjacent region that are required for designer sensor NLRs to activate RGA4-dependent immunity in rice.

In this study, we contrived four RGA5-HMA mutants based on the crystal structures of AVR-PikD, the RGA5-HMA/AVR1-CO39 complex and the Pikp-HMA/AVR-PikD complex. Intriguingly, all the mutants gained binding affinity to AVR-PikD, but only RGA5^HMA5^ harboring the RGA5-HMA5 ID, activated RGA4-dependent cell death of rice protoplasts and conferred resistance in transgenic rice plants to the *M. oryzae* strains expressing AVR-PikD. We identified three interfaces bound by AVR-PikD in the C-terminus of RGA5^HMA5^, two of which are located within the HMA ID and the other carrying a Lys-rich stretch tails the HMA ID. Notably, the RGA5 mutants having one or two of the three engineered interfaces failed to activate RGA4-dependent cell death in rice protoplasts. Further, the positively charged Lys residues in the Lys-rich stretch interface are essential to the derepression of RGA4 by RGA5^HMA5^. Altogether, this study demonstrates that synthetic sensor NLRs require a concerted action of multiple interfaces within and outside IDs for the effector-binding and receptor-activation functions, and has implications for structure-guided rational designing of sensor NLR receptors. This study also represents a significant advance towards designing NLR receptors with distinct specificity of recognition, which can be deployed for breeding multiline cultivars to help prevent the erosion of host resistance^20^.

## Results

### Designer NLR receptors carrying a single engineered effector-binding interface within the RGA5-HMA bind to the noncorresponding MAX effector AVR-PikD but fail to trigger RGA4-dependent cell death in rice protoplasts

Previous studies have revealed that the Pik1-HMA and the RGA5-HMA domains are structurally similar, comprising a four-stranded antiparallel β-sheet and two α-helices packed in an α/β sandwich model^10, 13^. RGA5-HMA domain interacts mainly with the β2 strand of AVR1-CO39 via the α1 helix-β2 strand interface^13^ (Fig. 1a). By engineering this interface along with the Lys-rich stretch located immediately after the HMA domain, we generated a designer NLR receptor RGA5^HMA2^ that confers resistance in transgenic rice plants to the *M. oryzae* strains expressing the noncorresponding MAX effector AVR-Pib^17^. Therefore, we reasoned whether this interface could be resurfaced to generate an RGA5-HMA mutant capable of binding to another noncorresponding effector AVR-PikD. Structural superimposition of AVR-PikD with the complex of AVR1-CO39/RGA5-HMA suggested that the M1016V mutation in RGA5-HMA may form hydrophobic interactions with A67 and G68 in AVR-PikD, and reduces steric hindrance at the interface between RGA5-HMA and AVR-PikD (Fig. 1a). The G1009D and S1027V mutations were adopted to block the interaction with AVR1-CO39 based on the previously contrived designer NLR RGA5^HMA2 17^. We thus generated a mutant of RGA5-HMA carrying the G1009D, M1016V and S1027V mutations, named RGA5-HMA3. A previous study reported that the β2-β3-β4 sheet within Pik-HMA is the interface for interaction with AVR-Pik^16^. E230 in the β3 strand of Pikp-HMA is one of the key residues interacting with H46 of AVR-PikD or N46 of other AVR-Pik effectors by a salt bridge or hydrogen bond, which corresponds to V1039 in RGA5-HMA that may decrease the interaction with AVR-PikD (Fig. 1b, d)^10, 21^. Further comparison of the interfaces in the structures of RGA5-HMA, Pikp-HMA, Pikm-HMA and the complexes of Pik-HMA bound to different AVR-Pik effectors suggested that E1070 immediately after the β4 strand of RGA5-HMA corresponds to the N261 residue of Pikp-HMA, which caused the “looping out” of this region of Pikp1-HMA, thereby decreasing the binding affinity as compared with Pikm1-HMA that lacks the N residue (Fig. 1c). We then generated another RGA5-HMA variant carrying the V1039E substitution and the E1070 deletion, named RGA5-HMA4, for recognizing AVR-PikD (Fig. 1d).

**Fig. 1.**
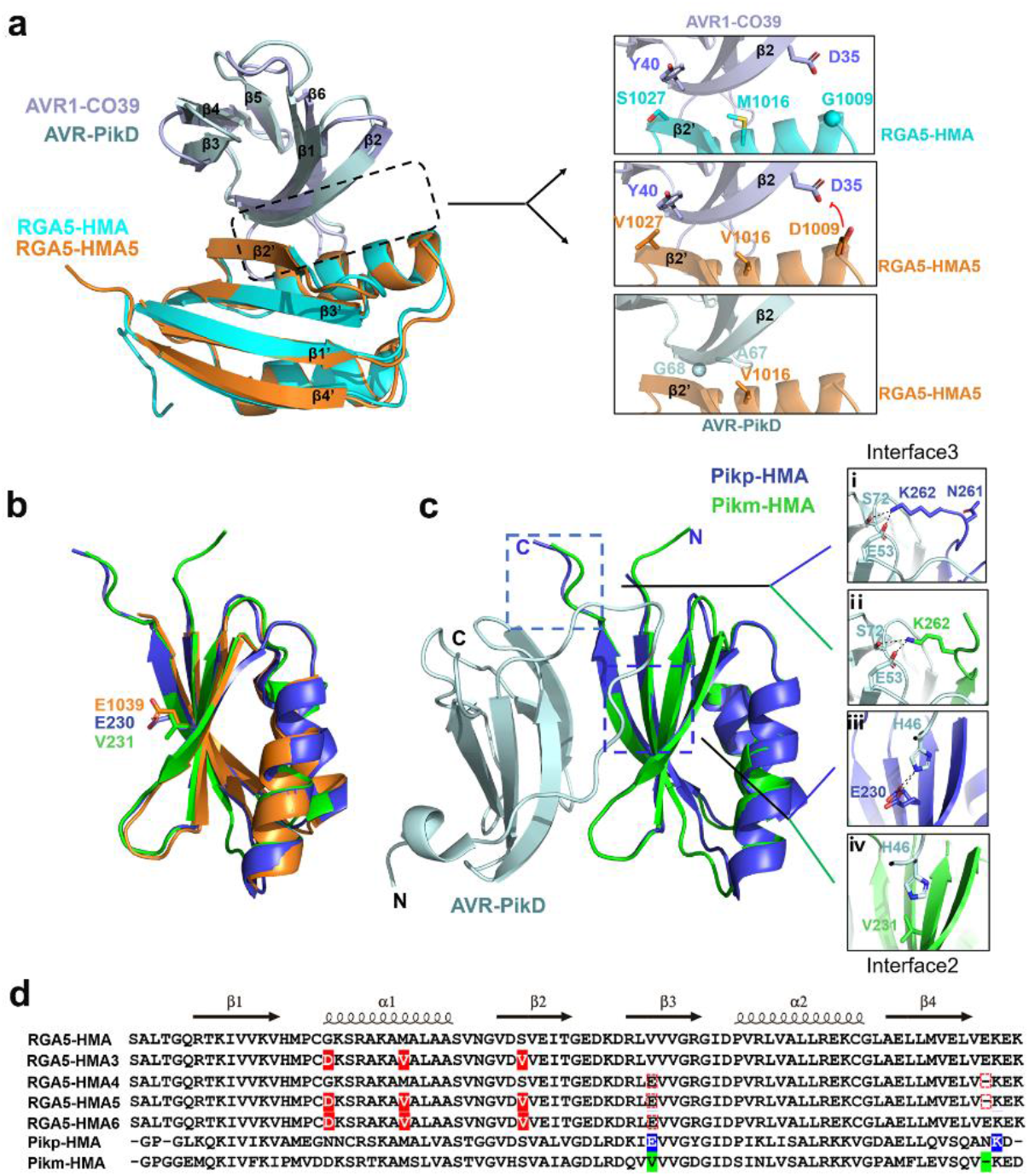
Designing RGA5-HMA mutants capable of recognizing the noncorresponding effector AVR-PikD. **a** The residues involved in the binding interface in the complexes of AVR1-CO39/RGA5-HMA and AVR-PikD/ RGA5-HMA5 were predicted by the structural superposition of the HMAs and the effectors. **b** The structural superposition of Pikp1- (PDB: 6G10), Pikm1- (PDB: 6FU9) and RGA5-HMA5 HMA domains. E1039 in RGA5-HMA5 labeled in orange corresponds to E230 in Pikp (blue) and V231 in Pikm (green) HMAs. **c** The two AVR-PikD-binding interfaces in Pikp1 or Pikm1 HMA domain. **d** Amino acid sequence alignment of RGA5-HMA mutants with RGA5-HMA and Pikp/Pikm-HMA. Labeled in the red box are modified residues in the RGA5-HMA mutants corresponding to the residues in Pikp1/Pikm HMA that were labeled in blue and green. Mutated residues in RGA5-HMA blocking the AVR-Pia binding are indicated in red shade. In RGA5-HMA mutants, E1070 was deleted, and V1039 was mutated into E, respectively. Secondary structural features of the HMA domains are shown above the alignment.

To test whether RGA5-HMA3 and RGA5-HMA4 are able to bind to AVR-PikD, we first performed the yeast two-hybrid (Y2H) assay. As shown in Fig. 2a, both RGA5-HMA3 and RGA5-HMA4 interacted with AVR-PiKD in the yeast cells. To further confirm the interactions in plant cells, we replaced the wild type RGA5-HMA with RGA5-HMA3 and RGA5-HMA4, generating two designer NLRs called RGA5^HMA3^ and RGA5^HMA4^. Co-IP assays showed that when coexpressed in *N. benthamiana*, both the HA-tagged RGA5^HMA3^ and RGA5^HMA4^ were coimmunoprecipitated with GFP-tagged AVR-PikD. In contrast, RGA5 was not coimmunoprecipitated with AVR-PikD (Fig. 2b). These results indicated that RGA5^HMA3^ and RGA5^HMA4^ could bind to the noncorresponding MAX effector AVR-PikD.

**Fig. 2.**
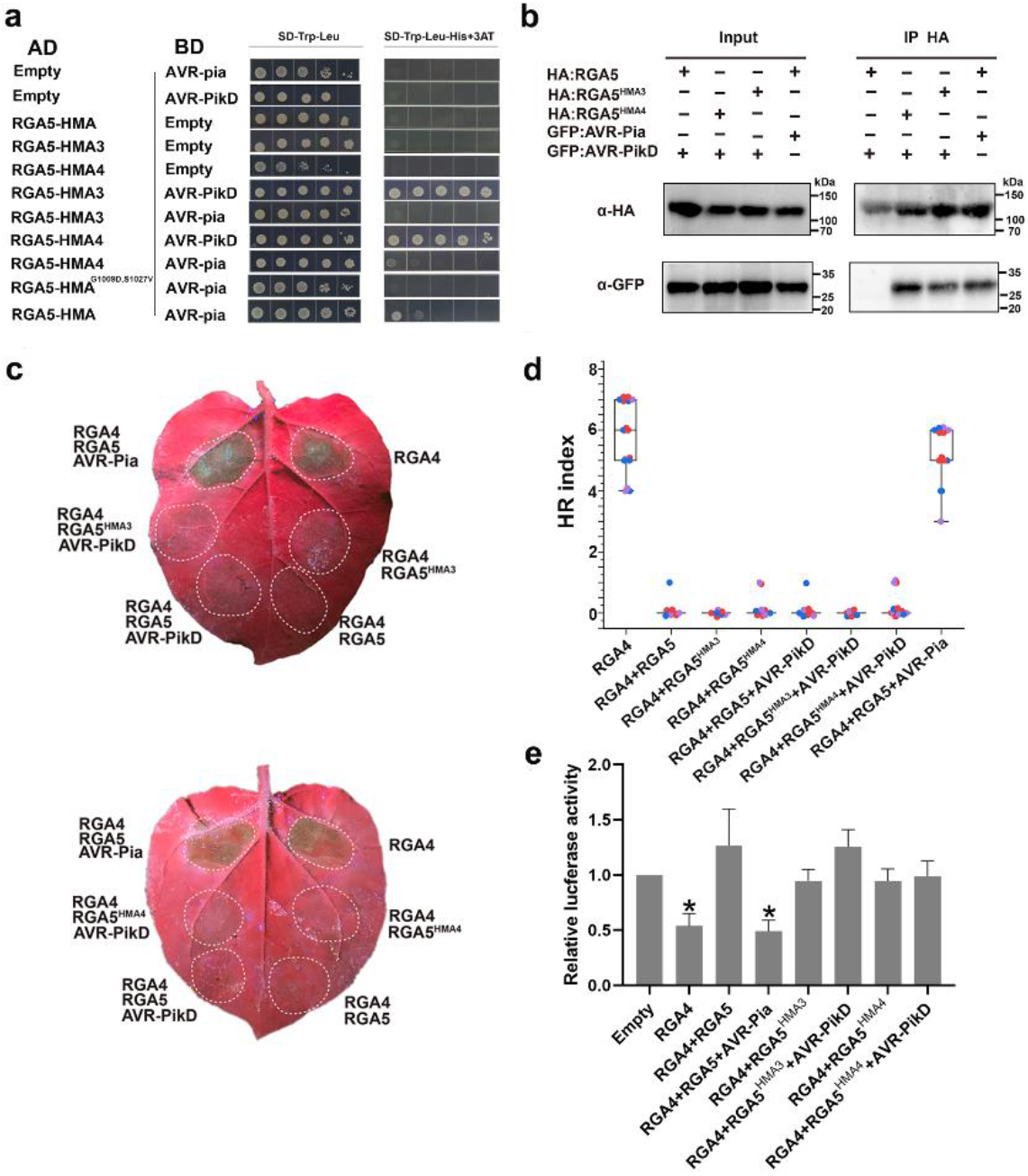
Functional analysis of RGA5^HMA3^ and RGA5^HMA4^ in the *N. benthamiana* leaves and rice protoplasts. The interaction of RGA5-HMA3 or RGA5-HMA4 with AVR-PikD was verified by Y2H **(a)** and Co-IP in *N. benthamiana* **(b)**. **c** Images of the *N. benthamiana* leaves coinfiltrated with RGA5^HMA3^ or RGA5^HMA4^, and RGA4, AVR-PikD and silencing suppressor p19. Leaves at three days post infiltrations were photographed under the UV light. **d** HR index of different combinations in representative pictures **(c)** was scored as previously reported^17^. Twenty biological replicates were used in each group. Three independent groups in different colors were labeled in box plots. Differences among the samples were assessed by Tukey’s HSD test (p<0.01). **(e)** The LUC activity in rice protoplasts cotransfected with different vector combinations. RGA4 was set as the positive control, and empty vectors served as the negative control. Significant differences with empty vector samples are labeled with an asterisk and assessed by Dunnett’s HSD test (*p*<0.05). The assays were repeated three independent times.

We further tested whether RGA5^HMA3^ and RGA5^HMA4^ could trigger RGA4-dependent cell death upon recognizing AVR-PikD in the *N. benthamiana* leaves and rice protoplasts. As shown in Fig. 2c and d, the cell death was not visible in either *Agrobacterium tumefaciens*-mediated coexpression of AVR-PikD and RGA4 with RGA5^HMA3^ or with RGA5^HMA4^ in *N. benthamiana* or cotransfection in the rice protoplasts. As a control, RGA4-dependent cell death was induced by the combination of AVR-Pia with RGA5 both in the *N. benthamiana* leaves and in the rice protoplasts (Fig. 2c, d). These results suggested that the binding of AVR-PikD by RGA5^HMA3^ or RGA5^HMA4^ is insufficient to trigger RGA4-dependent cell death, thus requiring further optimization of residues implicated in the receptor activation step.

### RGA5^HMA5^ carrying the mutations of RGA5^HMA3^ and RGA5^HMA4^ can trigger RGA4-dependent plant cell death upon the perception of AVR-PikD

Previous studies have revealed that RGA5-HMA recognizes AVR-CO39 with a distinct interface from that in Pik1-HMA for binding to AvrPikD^10, 13^ and that RGA4 is derepressed to activate plant cell death upon the recognition of AVR-Pia by RGA5^8, 14^. Since RGA5^HMA3^ and RGA5^HMA4^ carried a single engineered interface for binding to AVR-PikD and failed to trigger RGA4-mediated cell death, we reasoned that a functional designer RGA5 might require both an effector-binding interface and RGA4 derepression motifs within the HMA domain and its adjacent regions. Therefore, we created RGA5-HMA5 by combining the mutations present in RGA5-HMA3 and RGA5-HMA4. Meanwhile, we determined the crystal structure of RGA5-HMA5 and confirmed it was similar to that of the wild type RGA5-HMA (RMSD = 0.6 Å) (Fig. 1b and Supplementary Fig. 1). Y2H and MBP pull-down assays showed that RGA5-HMA5 specifically interacted with AVR-PikD but not with AVR-Pia, while RGA5-HMA interacted with AVR-Pia but not with AVR-PikD (Fig. 3a, b). Microscale thermophoresis (MST) analyses revealed that RGA5-HMA5 binds to AVR-PikD with a dissociation constant (*Kd*) of 12 μM, while RGA5-HMA binds to AVR-Pia with a *Kd* of 35 μM (Fig. 3c), indicating that RGA5-HMA5 has a higher binding affinity for AVR-PikD. We then generated RGA5^HMA5^ by replacing the wild type RGA5-HMA with RGA5-HMA5, and performed Co-IP in *N. benthamiana* by transiently coexpressing HA-RGA5^HMA5^/GFP-AVR-Pia, HA-RGA5^HMA5^/GFP-AVR-PikD, HA-RGA5/GFP-AVR-Pia or HA-RGA5/GFP-AVR-PikD. As shown in Fig. 3d, HA-RGA5^HMA5^ but not HA-RGA5 was coimmunoprecipitated with GFP-AVR-PikD. These results indicate that RGA5-HMA5 can specifically interact with AVR-PikD *in vivo* and *in vitro* (Supplementary Table 1).

**Fig.3.**
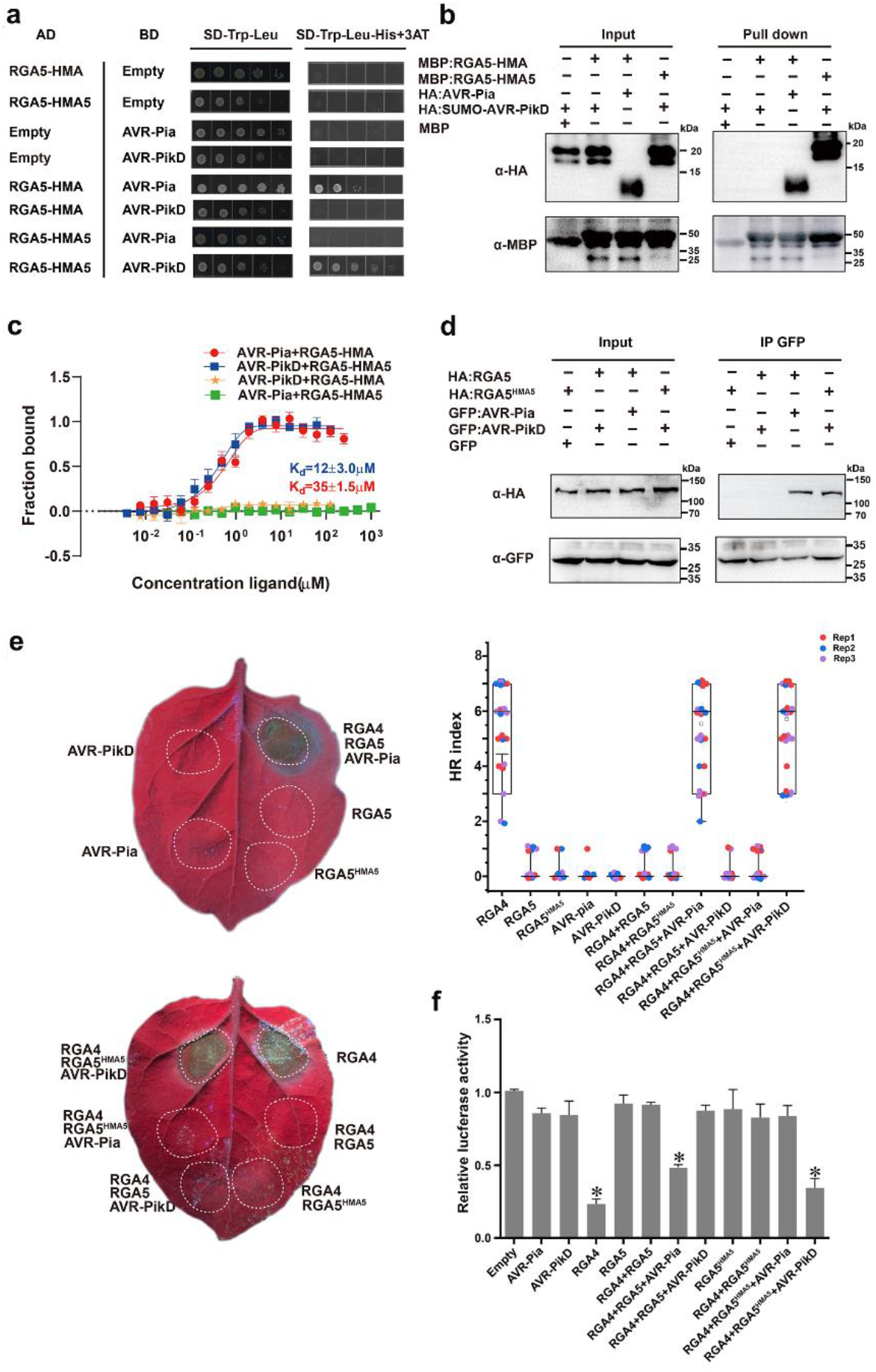
The designer NLR receptor RGA5^HMA5^ activates RGA4-dependent plant cell death upon recognizing the noncorresponding MAX effector AVR-PikD. **a** Y2H assays show the interaction of RGA5-HMA5 with AVR-PikD but not with AVR-Pia. **b** Pull-down assays show specific interaction of HA-AVR-PikD with MBP-RGA5-HMA5 but not with MBP-RGA5-HMA. HA-AVR-PikD, HA-AVR-Pia, MBP-RGA5-HMA5 and MBP-RGA5-HMA proteins were individually expressed in *E. coli*. Fusion proteins in different combinations were visualized by immunoblotting with the anti-HA and anti-MBP antibodies. **c** MST analyses show the dissociation constants (*Kd*) between the RGA5-HMA or RGA5-HMA5 domain and the effectors. The *Kd* and error bars were calculated from data from three independent biological replicates. **d** Coimmunoprecipitation assays showing the interaction of RGA5^HMA5^ with AVR-PikD in *N.benthamiana*. Co-IP proteins were detected by using anti-HA and anti-GFP antibodies, respectively. **e** Representative leaf images showing specific cell death in the *N. benthamiana* leaves after coinfiltration of the *A. tumefaciens* strains carrying RGA5^HMA5^ (fused with HA) with RGA4 (fused with Flag) and AVR-PikD (fused with GFP), or RGA5 (fused with HA) with RGA4 (fused with Flag) and AVR-Pia or RGA4. Images were taken three days after the infiltrations under the UV light. HR index was scored based on representative pictures as previously reported (Liu et al., 2021). Twenty biological replicates were used in each group. Three independent groups in different colors were labeled in box plots. Differences among the samples were assessed by Tukey’s HSD test (p<0.01). **f** The LUC activity in rice protoplasts after transfection with different vector combinations. RGA4 was set as the positive control, and the empty vectors were served as the negative control. Significant differences with empty vector samples are labeled with an asterisk and assessed by Dunnett’s HSD test (*p*<0.01). The assays were repeated three times with similar results.

**Table 1.**
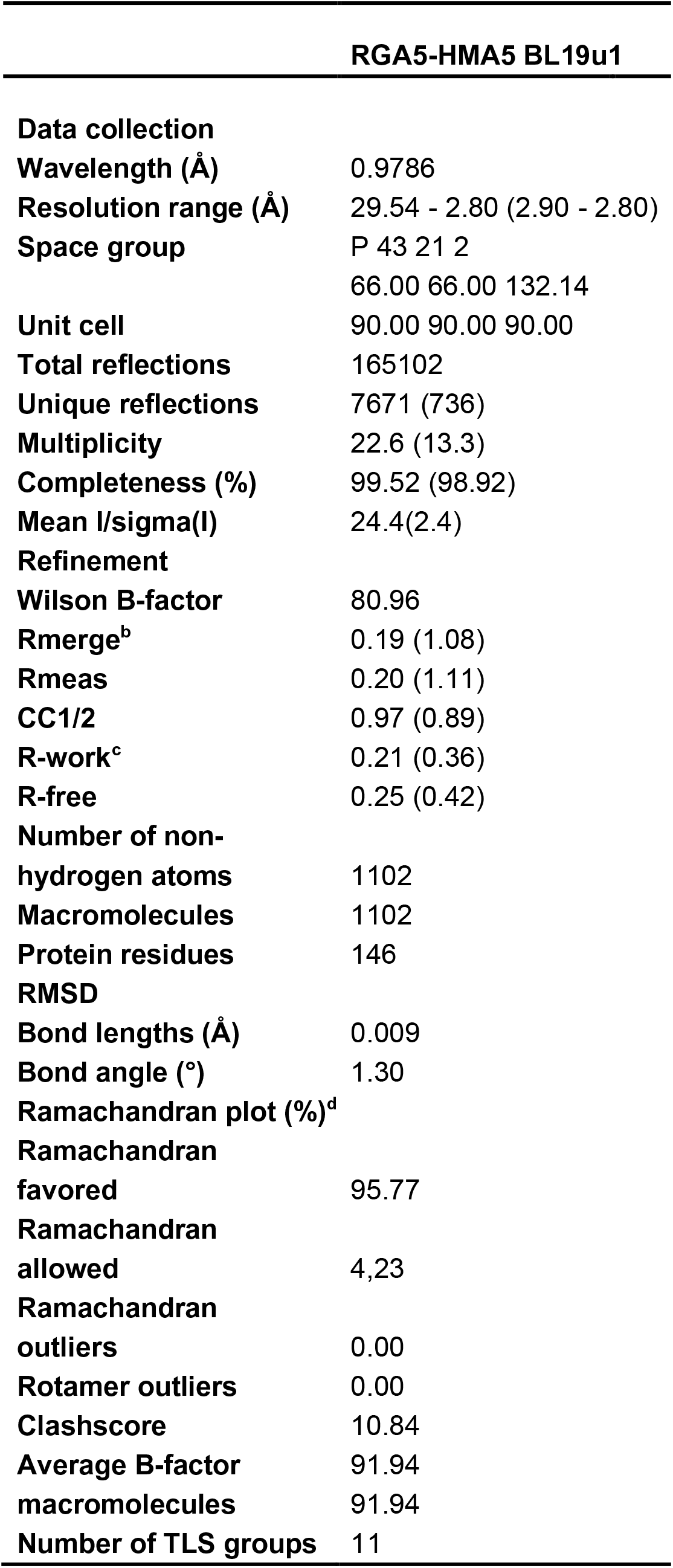
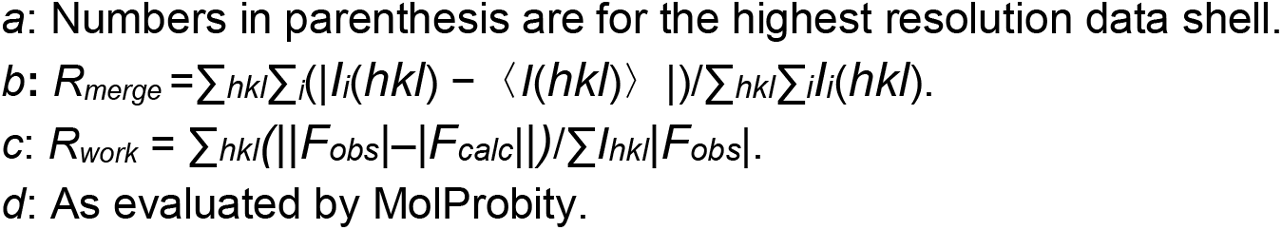
Data collection and refinement statistics.

| **RGA5-HMA5 BL19u1** |  |
| --- | --- |
| **Data collection** |  |
| **Wavelength (Å)** | 0.9786 |
| **Resolution range (Å)** | 29.54 - 2.80 (2.90 - 2.80) |
| **Space group** | P 43 21 2 |
|  | 66.00 66.00 132.14 |
| **Unit cell** | 90.00 90.00 90.00 |
| **Total reflections** | 165102 |
| **Unique reflections** | 7671 (736) |
| **Multiplicity** | 22.6 (13.3) |
| **Completeness (%)** | 99.52 (98.92) |
| **Mean I/sigma(I)** | 24.4(2.4) |
| **Refinement** |  |
| **Wilson B-factor** | 80.96 |
| **Rmerge<sup>b</sup>** | 0.19 (1.08) |
| **Rmeas** | 0.20 (1.11) |
| **CC1/2** | 0.97 (0.89) |
| **R-work<sup>c</sup>** | 0.21 (0.36) |
| **R-free** | 0.25 (0.42) |
| **Number of non-hydrogen atoms** | 1102 |
| **Macromolecules** | 1102 |
| **Protein residues** | 146 |
| **RMSD** |  |
| **Bond lengths (Å)** | 0.009 |
| **Bond angle (°)** | 1.30 |
| **Ramachandran plot (%)<sup>d</sup>** |  |
| **Ramachandran favored** | 95.77 |
| **Ramachandran allowed** | 4.23 |
| **Ramachandran outliers** | 0.00 |
| **Rotamer outliers** | 0.00 |
| **Clashscore** | 10.84 |
| **Average B-factor macromolecules** | 91.94 |
| **Number of TLS groups** | 11 |

a: Numbers in parenthesis are for the highest resolution data shell.
b: $R_{merge} = \sum_{hkl} \sum_i (|I_i(hkl) - \langle I(hkl) \rangle|) / \sum_{hkl} \sum_i I_i(hkl)$ .
c: $R_{work} = \sum_{hkl} (||F_{obs}| - |F_{calc}||) / \sum_{hkl} |F_{obs}|$ .
d: As evaluated by MolProbity.

To verify whether RGA5^HMA5^ is able to activate RGA4-dependent plant cell death upon recognizing AVR-PikD, the RGA4/RGA5^HMA5^ pair was first transiently coexpressed with AVR-PikD in the *N. benthamiana* leaves. As shown in Fig. 3e, the coexpression induced cell death similar to RGA4/RGA5 with AVR-Pia. In contrast, cell death was not visible when RGA4/RGA5 with AVR-PikD, RGA4/RGA5^HMA5^ with AVR-Pia, RGA4/RGA5, RGA4/RGA5^HMA5^, AVR-Pia or AVR-PikD were infiltrated in the *N. benthamiana* leaves (Fig. 3e) although the proteins were expressed in the leaves (Supplementary Fig. 2). We further measured the luciferase reporter activity in rice protoplasts after expressing the different combinations of proteins. As shown in Figure 3F, the coexpression of RGA5^HMA5^/RGA4 with AVR-PikD significantly reduced the luciferase reporter activity, similar to that of RGA5/RGA4 with AVR-Pia. However, such a reduction in the luciferase reporter activity was not detected by the coexpression of RGA4/RGA5 with AVR-PikD, and RGA4/RGA5^HMA5^ with or without AVR-Pia (Fig. 3f), and RGA4/RGA5m1 or RGA4/RGA5m1m2 with AVR-PikD (Supplementary Fig. 3). In addition, the expression of RGA5^HMA5^ or AVR-PikD alone in rice protoplasts could not cause an apparent immune response, indicating that RGA5^HMA5^ had no cell death-inducing activity but still retained the ability to repress the activity of the RGA4-induced cell death (Fig. 3f). Altogether, these results indicate that RGA5^HMA5^ gained the AVR-PikD recognition specificity *in planta*, thereby inducing RGA4-dependent cell death, consistent with the observations in the *N. benthamiana* leaves (Supplementary Table 1).

### Transgenic rice expressing RGA5^HMA5^ and RGA4 confers complete resistance to the blast fungus expressing AVR-PikD

The above results showed that only RGA5^HMA5^ among the three designer RGA5 mutants could cause RGA4-mediated cell death in rice protoplasts upon recognition of AVR-PikD (Fig. 2e, 3f). We thus generated five independent transgenic rice lines by cotransforming *RGA4* and *RGA5^HMA5^* into Nipponbare, a rice cultivar lacking *Pia* and *Pik*, and tested whether their T1 generation lines resist infection by the *M. oryzae* strains expressing *AVR-PikD*. The T1 lines of *RGA4/RGA5*^17^, two monogenic lines IRBLa (expressing only *Pia*) and IRBLk (expressing only *Pik*) of Lijiangxintuanheigu (LTH)^22^, and Nipponbare were used as controls. The *M. oryzae* wild-type strain DG7 lacking functional *AVR-Pia* and *AVR-PikD* and its transformants expressing *AVR-Pia* or *AVR-PikD* were used to infect the control and transgenic lines. As expected, the monogenic lines IRBLa and IRBLk were resistant to the DG7 transformants expressing *AVR-Pia* and *AVR-PikD*, respectively, while Nipponbare was susceptible to DG7 and its transformants expressing *AVR-Pia* or *AVR-PikD* (Fig. 4), confirming that the *M. oryzae* strains used in the assay were reliable. We then inoculated the transgenic rice lines expressing *RGA4/RGA5^HMA5^* or *RGA4/RGA5* by wound inoculation (Liu et al., 2021). As shown in Fig. 4, small resistant lesions were formed in the *RGA4/RGA5^HMA5^* transgenic lines after inoculation with the DG7 transformants expressing *AVR-PikD* but not with the wild-type strain DG7 or the transformants thereof carrying *AVR-Pia* (Fig. 4a, b). Meanwhile, the transgenic rice lines expressing *RGA4/RGA5* were resistant only to the transformants expressing *AVR-Pia*, but not to DG7 or the transformants expressing *AVR-PikD* (Fig. 4a, b). We further estimated *in planta* biomass of *M. oryzae* in the inoculated rice lines, verifying that the DG7 transformants expressing *AVR-PikD* were significantly limited in proliferation in the *RGA4/RGA5^HMA5^* transgenic lines and IRBLk but not in the other rice lines (Fig. 4c). In addition, a qPCR analysis confirmed that the NLR gene pairs and the effector genes were correctly expressed in the transgenic rice lines and *M. oryzae* during infection (Supplementary Fig. 4). Altogether, the above results demonstrated that the designer NLR receptor gene *RGA5^HMA5^* coexpressed with *RGA4* in transgenic rice plants could confer specific resistance to the *M. oryzae* strains expressing *AVR-PikD*, mimicking Pik/AVR-PikD-mediated resistance.

**Fig. 4.**
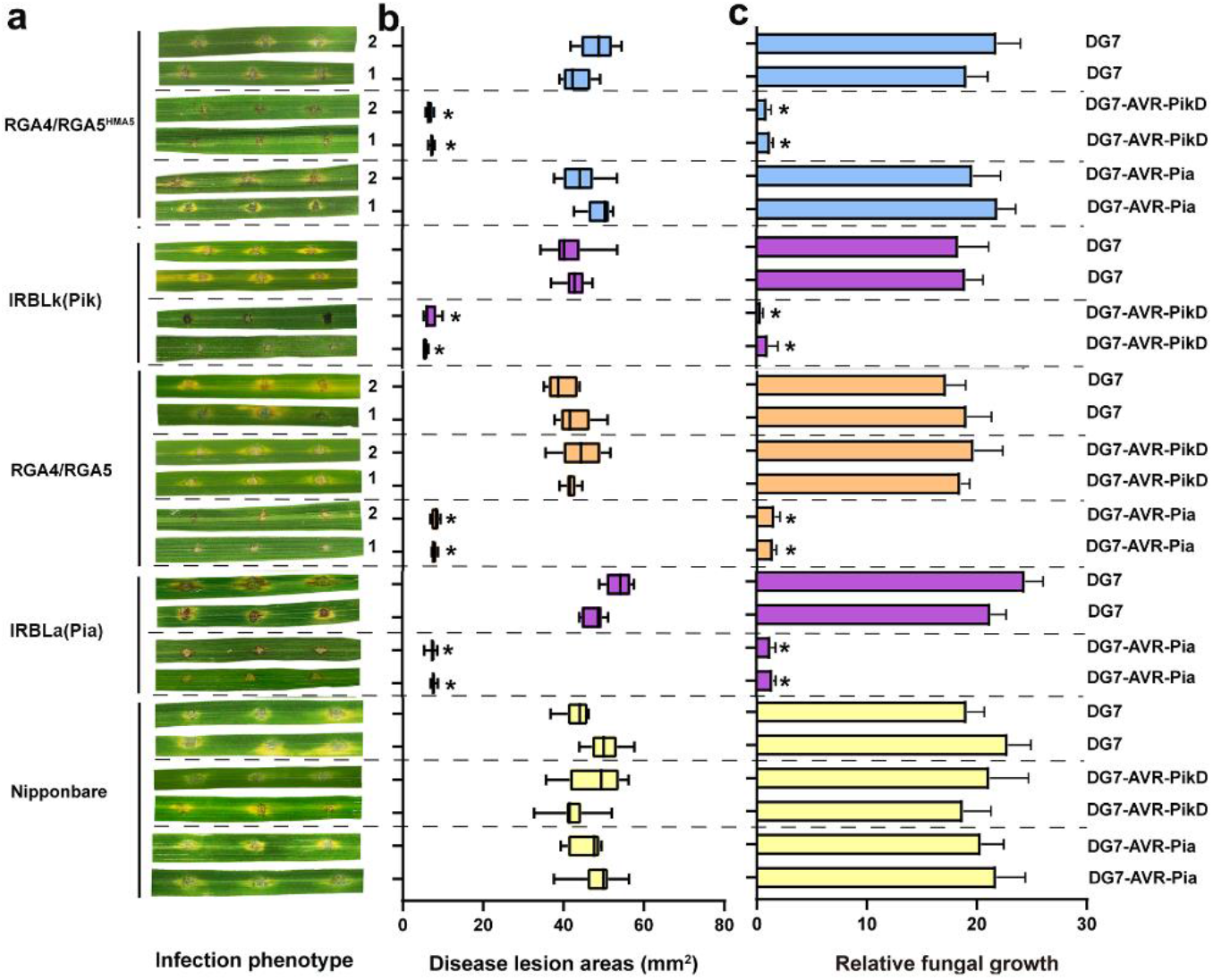
The transgenic rice lines expressing *RGA5^HMA5^*/*RGA4* confer specific resistance to the *M. oryzae* strains expressing the noncorresponding effector AVR-PikD. **a** Images showing disease reactions (resistant or susceptible) on the leaves of Nipponbare and its transgenic lines expressing *RGA4/RGA5^HMA5^* or *RGA4/RGA5*, and the LTH monogenic lines IRBLk (expressing *Pik*) and IRBLa (expressing *Pia*) following the inoculation with the *M. oryzae* DG7 strain expressing AVR-PikD or AVR-Pia. The wild-type strain DG7 and its transformant expressing *AVR-PikD* or *AVR-Pia* caused susceptible reactions on Nipponbare. The infection assays were performed in triplicate. Conidial suspensions of *M. oryzae* for all the inoculations were adjusted at a concentration of 10^5^/ml. Representative photos of the inoculated leaves from two independent lines were taken four days after inoculation. **b** box-and-whisker plots showing statistics on the sizes of lesions formed as described in (a). Lesion areas were measured by ImageJ. **c** Bar graphs showing the biomass of *M. oryzae* in the infected rice leaves as described in (a). The fungal biomass was quantified by measuring the expression levels of *MoPot*2 in relation to the rice ubiquitin gene. Values are means with standard deviations of nine independent biological replications from three independent rice lines. Significance analysis compared to Nipponbare is labeled with an asterisk and performed with Student’s t-test (*p* <0.05).

### RGA5^HMA5^ harbors multiple AVR-PikD-binding interfaces, including the lysine-rich stretch tailing the HMA ID

To understand how RGA5^HMA5^ triggers RGA4-dependent plant cell death, we identified peptides bound by AVR-PikD at the C-terminus of RGA5^HMA5^ (997-1116 aa), including the RGA5-HMA5 domain, by the hydrogen/deuterium exchange coupled with mass spectrometry (HDX-MS). As shown in Fig. 5a, there were six peptides in the RGA5^HMA5^ C-terminus bound by AVR-PikD. As expected, two peptides (1010-1018 aa and 1021-1026 aa) located within the interface of RGA5-HMA binding to AVR1-CO39 and Pik-HMA/AVR-Pia, and one peptide (1030-1039 aa) corresponded to the interface of Pik1-HMA binding to AVR-PikD (Fig. 1a, b)^13, 21, 23^. In the interface corresponding to the Pik1-HMA interface binding to AVR-Pik, E1039 of RGA5-HMA5 forms salt bridges with the side chains of H46 of AVR-PikD (Fig. 1c). Notably, two peptides (1068-1095 aa and 1109-1116 aa) identified were from the C-terminal Lys-rich tail immediately following RGA5-HMA5 (Fig. 5a). Within this interface, the forward shift of K1070 resulted from the front E deletion in RGA5-HMA5 was designed to mimic K262 of Pikm-HMA (Fig. 1c), which interacts with E53 of AVR-PikD by a salt bridge^21^. To verify the significance of K1070 and V1039E in derepressing the RGA5^HMA5^/RGA4 complex, we generated the E53A and H46A mutations within AVR-PikD, and the mutants of RGA5-HMA6 and RGA5^HMA6^, in which E1070 was kept as in RGA5. E53A but not H46A mutation in AVR-PikD abolished the interaction with RGA5-HMA5 (Fig. 5b). To our surprise, RGA5^HMA6^ failed to trigger RGA4-dependent cell death by AVR-PikD, although RGA5-HMA6 retained AVR-PikD binding capability (Fig. 5b, c), suggesting that the E1070 deletion with the forward shift of K1070 is crucial to the capability of RGA5^HMA5^ to derepress RGA4 for inducing the rice immunity. Previous studies reported that, in contrast to other MAX effectors, AVR-PikD has an N-terminal negatively charged patch consisting of Asp and Glu residues^10^ (Fig. 5a and Supplementary Fig. 5). Y2H assays showed that AVR-PikD without the N-terminal Loop (named AVR-PikD^ΔN^) failed to interact with Pikm-HMA, RGA5-HMA5, RGA5-HMA3 and RGA5-HMA4 (Fig. 5b). In addition, the N-terminal Loop alone was unable to interact with the C-terminal tail of RGA5-HMA5 (Fig. 5b). Furthermore, as mentioned above, two peptides bound by AVR-PikD were located within the C-terminal Lys-rich stretch tailing the HMA ID, which is absent from Pik1-HMA^8, 17^. In a previous study, we showed that substituting all the positively charged Lys residues with negatively charged Glu residues in the C-tail is essential to the interaction of the designer receptor RGA5^HMA2^ with the noncorresponding AVR-Pib. We thus generated RGA5-HMA5^K/E^, an RGA5-HMA5 mutant, by replacing all the Lys residues with the Glu residues except K1070 in the C-terminal tail located immediately after RGA5-HMA5. As shown in Fig. 5b and 5c, RGA5^HMA5K/E^ could interact with AVR-PikD but lost the capability to trigger RGA4-dependent cell death in rice protoplasts by AVR-PikD, suggesting that the C-terminal positively charged Lys residues are crucial to the capability of RGA5^HMA5^ to derepress RGA4 for inducing the rice immunity. In addition, RGA5-HMA5 had a peptide bound by AVR-PikD that was located at the α2 helix-loop5 (Fig. 5a). Altogether, the above results revealed that RGA5^HMA5^ harbors at least three interfaces for the interaction with AVR-PikD, including an interface within the C-terminal Lys-rich tail, in which the positively charged residues are crucial for RGA5^HMA5^ to derepress RGA4 for triggering the rice cell death.

**Fig. 5.**
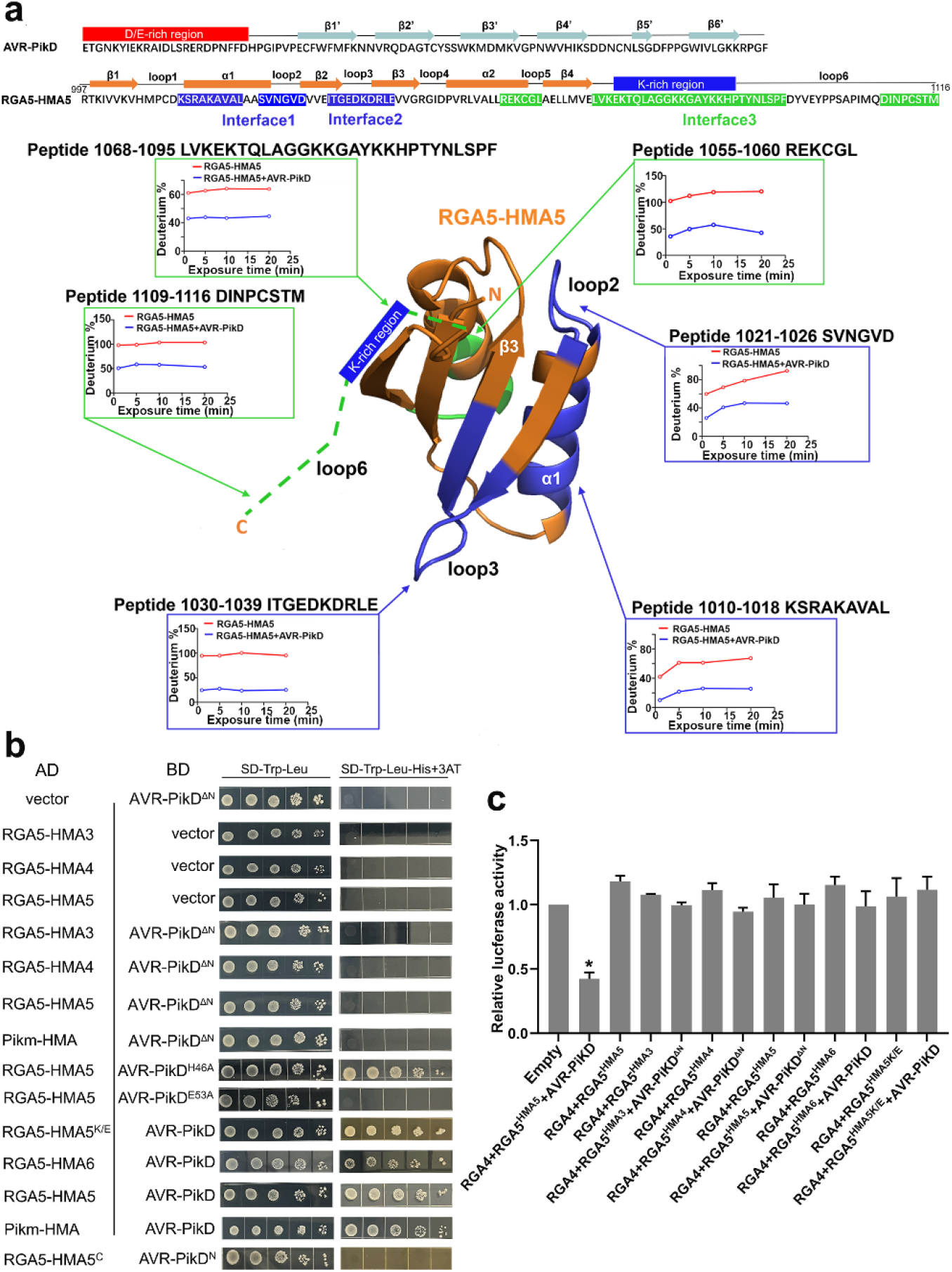
AVR-PikD-binding Interfaces of RGA5^HMA5^ identified by the epitope mapping based on hydrogen/deuterium exchange mass spectrometry (HDX-MS). **a** The C-terminus of RGA5^HMA5^ (997-1116 aa) was used to map peptides bound by AVR-PikD. The interaction interfaces are colored in blue and green on the top, and the complex model of RGA5-HMA5 bound to AVR-PikD is shown at the bottom. The green dashed lines indicate the C-terminal Lys-rich region after RGA5-HMA5. The graphs showed the deuterium percentage of HDX data over 20 min labeling for the C-terminal fragments of RGA5^HMA5^ corresponding to secondary structures in the absence (red) or presence (blue) of AVR-PikD. Significant reduction in the values of deuterium percentage indicates less exchange and more opportunities for binding to AVR-PikD. **b** Y2H assays show that the N-terminal loop of AVR-PikD is crucial to the interaction between the HMA domains and the effector. AVR-PikD^ΔN^ and RGA5-HMA5^K/E^ indicate the AVR-PikD mutant lacking the N-terminal loop and the RGA5-HMA5 mutant carrying the Glu substitutions of Lys residues in the C-terminal tail after the HMA domain, respectively. RGA5-HMA5^C^, the C-tail fragment after the HMA domain of RGA5^HMA5^; AVR-PikD^N^, the N terminal loop fragment of AVR-PikD. The others were as described in figure 1. **c** A bar graph shows that the LUC activity was significantly reduced in rice protoplasts after transfection with the RGA5^HMA5^/RGA4/AVR-PikD vector combination but not either by the RGA5^HMA6^/RGA4/AVR-PikD nor by the RGA5^HMA5K/E^/RGA4/AVR-PikD vector combination. Significant differences with empty vector samples are labeled with an asterisk and assessed by Dunnett’s HSD test (*p*<0.05). The assays were repeated three independent times.

## Discussion

Structure-guided rational engineering of NLRs is emerging as a promising approach to altering their recognition spectra and specificities ^24^. Several proof-of-concept studies have been recently reported on engineering NLRs and their IDs. The first designer NLR receptor was Pikp-1^NK-KE^, which was generated by engineering the Pikp-1 HMA ID, showing an expanded recognition spectrum against related AVR-Pik alleles^16^. Kourelis *et al*. contrived a series of Pikm-1 mutants called Pikobodies by replacing the Pikm1 HMA ID with nanobodies of fluorescent proteins, which could recognize antigen fluorescent proteins and trigger immunity in *N. benthamiana*^25^. Tamborski *et al*. created a mutant of wheat NLR receptor Sr33 capable of recognizing the avirulence effector AvrSr50 from stem rust by switching amino acid residues in the LRR domain with the AvrSr50-binding residues in Sr50^26^. However, the functionality of these designer NLRs remains to be verified in their host plants. We recently reported a designer NLR named RGA5^HMA2^ that conferred complete resistance in rice to the *M. oryzae* expressing the non-corresponding avirulence effector AVR-Pib^17^. RGA5^HMA2^ is the first example of designer NLRs conferring complete resistance in plants^24^. Here, we report yet another designer rice NLR receptor named RGA5^HMA5^, which conferred complete resistance in transgenic rice plants to the *M. oryzae* strains expressing the noncorresponding effector AVR-PikD. More importantly, we delimit and resurface the interfaces within and outside the HMA ID of RGA5, which not only impart *de novo* effector binding but also activate the NLR-mediated resistance. Our studies demonstrate that rationally engineering the HMA ID in RGA5 can generate a series of designer NLR receptors with distinct resistance profiles^17^, which will be useful for efficiently breeding multiline cultivars to maintain the durability of NLR-mediated resistance^20, 27^.

Wang et al. created a variant of RRS1 by adding a *Phytoplasma* effector SAP05-dependent degron domain to the C-terminus of RRS1, which elicited the hypersensitive response (HR) in *N. tabacum* leaves triggered by the effector but failed to confer full resistance in transgenic *Arabidopsis* to the *Phytoplasma* infection^19^. By engineering RGA5-HMA in accordance with Pikp1-HMA, Cesari *et al*. generated two designer NLR receptors, which shared the engineered β2β3β4 interface of RGA5-HMA and gained a high binding affinity to the non-corresponding effector AVR-PikD in addition to the corresponding effector AVR-Pia, but were unable to confer specific blast resistance in transgenic rice plants against *M. oryzae* strains expressing AVR-PikD^18^. Similarly, RGA5^HMA3^, RGA5^HMA4^ and RGA5^HMA6^ generated in this study, which carried a single or two engineered interfaces, also failed to trigger RGA4-dependent rice immunity. Compared to these designer RGA5 receptors unable to trigger RGA4-dependent rice immunity, RGA5^HMA5^ contains two engineered interfaces within RGA5-HMA and one interface in the Lys-rich stretch tailing the HMA ID (Supplementary Fig. 6), indicating that synthetic sensor NLR receptors may require multiple engineered interfaces to be active in host plants. Notably, we showed that replacing the five Lys residues with Glu residues in the C-terminal Lys-rich tail abolished the capability of RGA5^HMA5^ to induce RGA4-dependent cell death, indicating that the adjacent C-terminal tail functions not only as an interface for binding to AVR-PikD but also plays a regulatory role for RGA5^HMA5^ to derepress RGA4. Regulation of IDs by their adjacent domains may not be unique to RGA5. A previous study showed that DOM 4 and DOM 6 adjacent to the WRKY ID in RRS1 contribute to autoinhibition and activation of RRS1/RPS4 immune receptor complex^28^. Therefore, concurrent modification of IDs and their adjacent domains rather than ID alone may be a prerequisite to creating designer sensor NLR receptors.

We previously reported that RGA5^HMA2^ gained specific resistance to the *M. oryzae* strains expressing the noncorresponding MAX effector AVR-Pib but lost the inherent resistance to the *M. oryzae* strains expressing the corresponding MAX effector AVR-Pia^17^. Here again, RGA5^HMA5^ lost resistance to the *M. oryzae* strains with AVR-Pia, although it gained the capability to resist infection by the *M. oryzae* strains expressing the noncorresponding MAX effector AVR-PikD. In contrast, RGA5m1 and RGA5m1m2 retained the resistance of RGA5 to the *M. oryzae* strain carrying the corresponding AVR-Pia but did not gain the capability to resist infection by the *M. oryzae* strains carrying the noncorresponding effector AVR-PikD^18^. These studies raise a key question of whether molecular engineering of IDs in sensor NLR receptors can confer an expanded spectrum of resistance to pathogens expressing unrelated effectors. However, Maidment *et al*. recently generated two variants of Pikp-1 with expanded blast resistance profiles in transgenic rice plants^29^. Therefore, further investigation is required to optimize reported designer RGA5 receptors and determine the relationship between the interfaces and key residues thereof for binding to the corresponding and noncorresponding effectors.

*N. benthamiana* or *N. tabacum* has been widely adopted as a convenient heterologous system to assay plant immune responses, such as HR. However, recent studies showed that designer NLR receptors that gained the capability to recognize effectors and trigger HR in *N. benthamiana* or *N. tabacum* may not always enable complete resistance in host plants^18, 19^. These studies alert us that using the heterologous overexpression system to assess the synthetic NLRs seems insufficient. As such, there is a need for a homologous transient expression system to guide us on NLR gain-of-function engineering. We showed that RGA5-based designer NLRs were able to trigger RGA4-dependent cell death in rice protoplasts and confer complete blast resistance in transgenic rice, indicating that the cell death in rice protoplasts induced by RGA5-based designer NLRs is consistent with the intact plant blast resistance. Therefore, using homologous system-based bioassays prior to making transgenic plants may be a prerequisite to determining whether an engineered NLR gains recognition capacity to confer resistance.

In summary, we created a designer NLR receptor RGA5^HMA5^ that confers full resistance in transgenic rice plants to the blast fungus *M. oryzae* strains expressing the noncorresponding effector AVR-PikD. More importantly, we show that synthetic sensor NLR receptors require concurrent structure-guided engineering of multiple interfaces within and outside IDs. Notably, we found that the C terminal lysine-rich stretch tailing the HMA ID in RGA5^HMA5^ is an interface crucial to both recognizing the MAX effectors and activating RGA4-dependent rice immunity. We also suggest that utilizing homologous systems is important to assay designer NLR receptors. This study not only represents a significant advance towards structure-guided rational engineering of NLR receptors but also has implications for designing sensor NLRs by engineering their IDs.

## Methods

### Generation of constructs

*RGA5-HMA* (nucleotides 2944 to 3348) were obtained by gene synthesis (Genecreate, Wuhan). Point mutations were generated by the Quik Change Site-Directed Mutagenesis Kit (Transgen). For the *E. coli* protein expression, RGA5-HMA and RGA5-HMA5 with HA-MBP-tag were independently ligated to the pETMBP1a vector, and AVR-PikD with HA-sumo-tag and AVR-Pia with HA-tag ligated to pETSUMO1a and pHAT_2_, respectively. For the Y2H assay, the *RGA5-HMA* mutants and the effector genes (*AVR-Pia* and *AVR-PikD*) were separately cloned into pGADT7 (Clontech) and pGBKT7 (Clontech). For the transient expression assays in *N. benthamiana*, RGA5, RGA5^HMA5^ and RGA4 fused with HA or Flag, and AVR-Pia and AVR-PikD effectors with GFP tag were individually cloned into the pCAMBIA 1305 vector. For the assays using rice protoplasts, *RGA4*, *RGA5*, *RGA5^HMA5^*, *RGA5^HMA6^*, *RGA5^HMA5K/E^*, *LUC*, *AVR-Pia* and *AVR-PikD* were inserted into pUC19. For rice transformation, *RGA*5 and *RGA*5^HMA5^ and *RGA*4 under their native promoters were separately inserted into the vectors pCAMBIA1305 and pCAMBIA1300, respectively. For *M. oryzae* transformation, *AVR-Pia* and *AVR-PikD* with their native promoters were respectively cloned into pKN. The primers used to amplify the abovementioned genes are listed in Supplementary Table 1.

### *M. oryzae* strains and rice transformation

Routine maintaining and conidia production of the wild-type strain of *M. oryzae* DG7 and its transformants carrying *AVR-Pia* or *AVR-PikD* were performed as previously reported^30^. DG7 protoplasts were transformed with the linearized pKN vectors carrying *AVR-Pia* or *AVR-PikD*, and the resulted transformants were screened as reported^31^. Rice cultivar Nipponbare lacking *Pia* and *Pik* was transformed as described previously^32^.

### Infection assays of the *M. oryzae* strains on rice lines

Conidia of *M. oryzae* DG7 and its transformants were adjusted at a concentration of 10^5^ conidia per ml with sterilized water containing 0.025% Tween-20 and wound-inoculated on the leaves of Nipponbare, two Lijiangxintuanheigu monogenic lines IRBLa (with *Pia*) and IRBLk (with *Pik*), and the transgenic rice lines. The inoculated leaves were incubated as described previously^17^. Disease lesions formed on the leaves were scored four days post-inoculation and measured with ImageJ (https://imagej.net/). The infection assay of rice lines by *M. oryzae* was repeated three times at least.

### qPCR assays

Total RNA was extracted from the *M. oryzae*-infected rice leaves using the RNA extraction kit (Vazyme, Nanjing) and then reverse transcribed into cDNA with the HiScript 1st Strand cDNA Synthesis Kit (Vazyme, Nanjing). qPCR was performed by ABI Quantstudio 6 Flex PCR program (Thermo Fisher Scientific), with the actin genes in *M. oryzae* and rice as internal controls for normalizing the gene expressions. The assays were performed with three independent replicates. Primers used for the assays were listed in Supplementary Table S2.

### Yeast two-hybrid (Y2H) assays

*AVR-PikD* and *AVR-Pia* without the signal peptide sequence were cloned into the plasmid pGBKT7 as bait vectors, whereas *RGA5* and its HMA domain mutants into pGADT7 as prey vectors. The yeast strain Y2H *AH109* was cotransformed with the prey and corresponding bait vectors following the protocol provided by the Yeastmaker™ yeast transformation system (Clontech). The yeast transformants were grown on SD/-Trp/-Leu medium, and the interactions were assessed on SD/-Trp/-Leu/-His plates with X-α-gal.

### Cell death assays in *N. benthamiana* leaves and rice protoplasts

As described previously^33^, the transient protein expression in *N. benthamiana* was used as a heterologous system to assay plant cell death triggered by AVR-Pia or AVR-PikD along with RGA4 and RGA5 or its mutants. The *A. tumefaciens* GV3101 strains containing the different constructs were prepared and infiltrated into the *N. benthamiana* leaves as described^17^. After 48 h incubation in the dark, cell death around the infiltration sites was scored and photographed under the UV light ^34^.

Luciferase activity in rice protoplasts was used as an indicator of host cell death triggered by RGA4 and RGA5 or its mutants along with AVR-Pia or AVR-PikD. The activity was measured by using the luciferase assay system (Promega), which was performed 16 hours after transfection with the plasmid combinations containing AVR-Pia or AVR-PikD along with RGA4 and RGA5 or its mutants. Rice protoplasts were prepared from Nipponbare leaves as described previously^33^, and the plasmid combinations mixed with the empty and LUC plasmids were cotransfected into rice protoplasts via the polyethylene glycol method^35^. The assay was repeated in three independent experiments.

### Co-IP and Immunoblotting

The *N. benthamiana* leaves infiltrated with different *GV3101* strains as described in the cell death assay were ground into powder with liquid nitrogen and then homogenized with the extraction buffer (25 mM Tris-HCL pH 7.5, 1 mM EDTA, 150 mM NaCl, 10 % glycerol, 2 % polyvinylpolypyrolidone [PVPP], 5 mM DTT, 1x protease inhibitor, and 1 mM PMSF). The homogenates were centrifugated, and the supernatant was then applied to anti-GFP agarose beads (50 μl) at 4°C for 3 h. The beads were washed three times with the washing buffer (25 mM Tris-HCL pH 7.5, 1 mM EDTA, 150 mM NaCl, 10 % glycerol, 5 mM DTT, and 1x protease inhibitor). The proteins were separated by 10 % SDS-PAGE gels and detected by immunoblotting with the first anti-GFP or HA-tag antibodies (Sigma) and the second anti-rabbit IgG-peroxidase antibody (Sigma). Membranes were washed three times with TBST buffer (20 mM Tris, pH 8.0, 150 mM NaCl, 0.05 % Tween-20), and the immunoblot signals were visualized using the HRP chemiluminescent substrate (Millipore). Rubisco small submit stained by ponceau was set to verify equal protein loading as the control.

### Interaction assays by Pull-down and Microscale Thermophoresis

For the pull-down assay, MBP-RGA5-HMA5, MBP-RGA5-HMA, HA-AVR-Pia and HA-SUMO-AVR-PikD proteins were expressed in *E. coli* and purified as described previously^13^. Recombinant RGA5-HMA, RGA5-HMA5 with the effector proteins in a binding buffer (20 mM Tris, 150 mM NaCl, 5 mM DTT, 4 mM EDTA, pH 7.4, 1 % Triton X-100) were mixed and applied for Anti-MBP beads (50 μl), incubated with gentle rotation for 3 h at 4°C. Then the resin was washed five times with the binding buffer and boiled for 10 min. All the proteins were loaded onto 10% SDS-PAGE gel separation, then transferred onto the PVDF membrane (Millipore), and subsequently detected with the anti-MBP antibody.

For the MST assay, RGA5-HMA and RGA5-HMA5 were separately labeled with the fluorescent dye NT-647 from kit MO-L001 of Nano Temper. All the labeled proteins were dissolved in the buffer containing 20 mM PBS, 150 mM NaCl, 0.05 % (v/v) Tween 20, pH 7.4, and mixed with different concentrations of effectors (AVR-PikD or AVR-Pia). Finally, the Kd (the dissociation constant) values between the effector and the RGA5 HMAs were measured using Monolith NT.115 (NanoTemper Technologies) with 30 % LED power and fitted with Nano Temper Analysis Software (Version 1.5.41). The assays were repeated in three independent experiments.

### Crystallization, data collection and structure determination of RGA5-HMA5

For protein crystallization, RGA5-HMA5 (residues 982-1116) was expressed and purified and concentrated to 7 mg/ml as described previously^36^. Crystals of the RGA5 HMA mutant were produced by sitting drop vapor diffusion, which occurred after three days under the same condition as the wild type RGA5-HMA (0.2 M ammonium nitrate, 20 % [w/v] PEG 3350). The crystals were soaked into cryoprotectant containing 20 % (v/v) glycerol and flash cooling into liquid nitrogen. X-ray diffraction data were collected at the Shanghai Synchrotron Radiation facility by beamline BL19U. Data were processed using the HKL-2000 processing package^37^. The structure was solved by molecular replacement using Phaser with 5ZNE as a search model. The final structure was obtained by rebuilding using Coot^38^, and further refined using PHENIX with TLS restraints^39^. The detailed statistics on data collection and refinement are listed in Table 1.

### HDX-MS

RGA5-HMA5 and the complex of RGA5-HMA5 and AVR-PikD were diluted 10-fold in the deuterium labeling buffer (20 mM PBS, 7.4, 150 mM NaCl, 99.8% D2O) and incubated at 25 °C. The labeling reactions were stopped at special time points (1, 5, 10, and 20 min) by adding 50 μL of ice-cold quenching buffer (4 M guanidine hydrochloride, 200 mM citric acid, and 100 mM TCEP, pH 1.8). After adding 5 μL pepsin solution (1μM) to the reactions for 3 min digestion, the quenched samples were applied to Thermo-Dionex Ultimate 3000 HPLC system autosampler. The peptides were separated using a 40-min linear gradient acetonitrile-water (8 to 50% containing 0.1% formic acid) at a flow rate of 125 μL/min on a reverse-phase column (ACQUITY UPLC BEH C18, 1.7 μM) with Thermo-Dionex Ultimate 3000 HPLC system connected to a Thermo Scientific Q Exactive mass spectrometer. Mass spectra data were acquired in resolution mode over an m/z range of 50 to 2000. The captured peptides were identified with Proteome Discoverer (version PD1.4 from Thermo Fisher Scientific) and HDX-MS data were processed by HD Examiner.

## Acknowledgments

This work was supported by grants from the Natural Science Foundation of China (Grant No. 32030089), the National Rice Industry Program from Ministry of Agriculture and Rural Affairs (CARS-01-16), the IRT Program (Grant No. IRT1042) and the 111 Project (Grant No. B13006). The authors thank BL17U1 at Shanghai Synchrotron Research Facility (SSRF) beamline and BL19U1 at National Facility for Protein Science in Shanghai, Zhangjiang Laboratory, China, for providing technical support and assistance in data collection and analysis. We also thank Sheng Yang He at Duke University for the critical reading of the manuscript and Xiaolin Tian in Tsinghua University Branch of China National Center for Protein Sciences Beijing for technical help in the HDX-MS experiment.

## Author Contributions

Y.L.P. and J.L. designed research; X.Z., Y.L., G.Y., T.Z., X.W., M.M., and L.G. performed research; Y.L.P., J.L., X.Z., Y.L, G.Y., T.Z., X.W., M.M., D.W., L.G., V.B. and H.G. analyzed data; and Y.L.P., J.L., X.Z., V.B. H.G. and Y.L. wrote the paper.

## Data deposition

The coordinates and structure factors have been deposited in the Protein Data Bank with accession code 7DVG (HMA5).

## Supplementary information

**Fig. S1.**
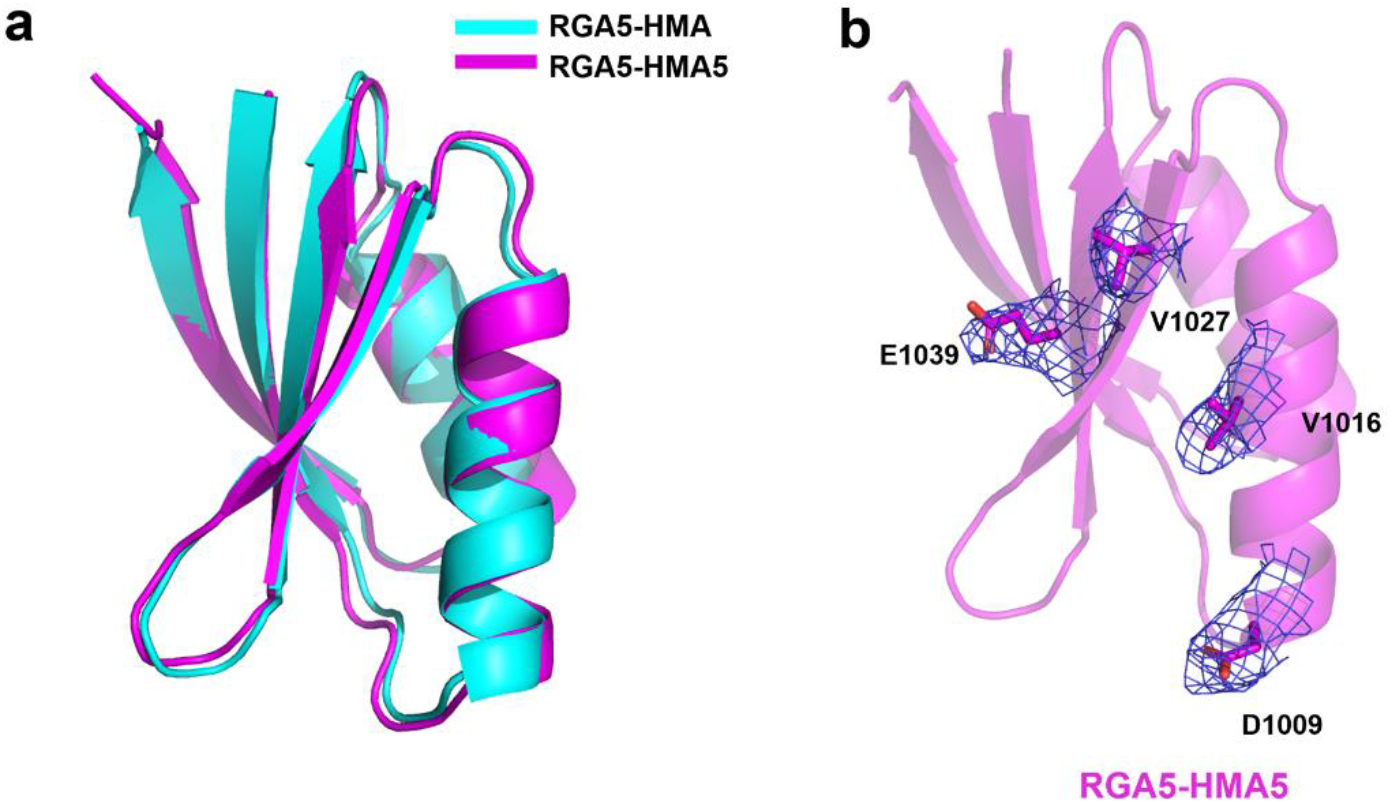
Superposition structure of RGA5-HMA and RGA5-HMA5 (A) and the modified residues of HMA5 domain were shown in blue mesh with the *2mFo-Fc* electron density map contoured at 1.0 σ (B).

**Fig. S2.**
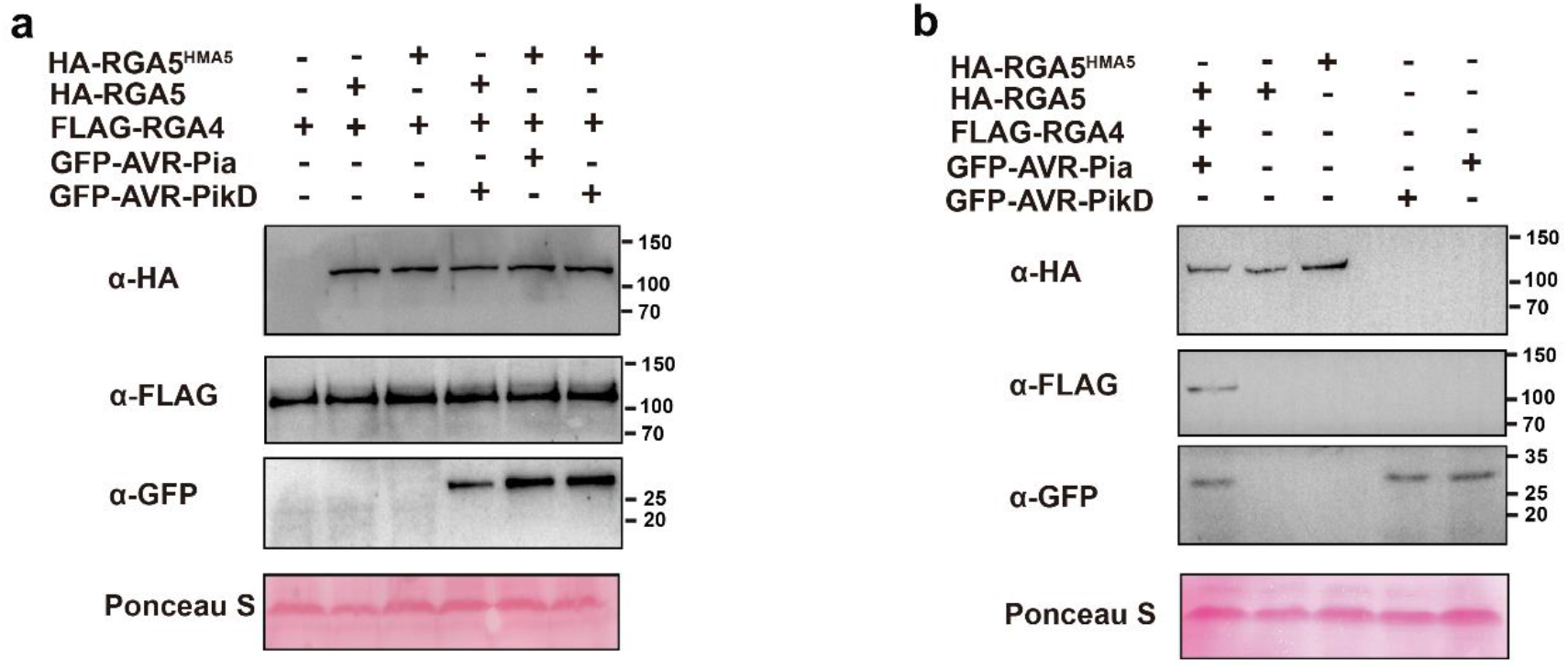
Expression of the fusion protein in different combinations visualized in Fig.3 by immunoblotting. Rubisco small submit stained by ponceau was used to verify equal protein loading.

**Fig. S3.**
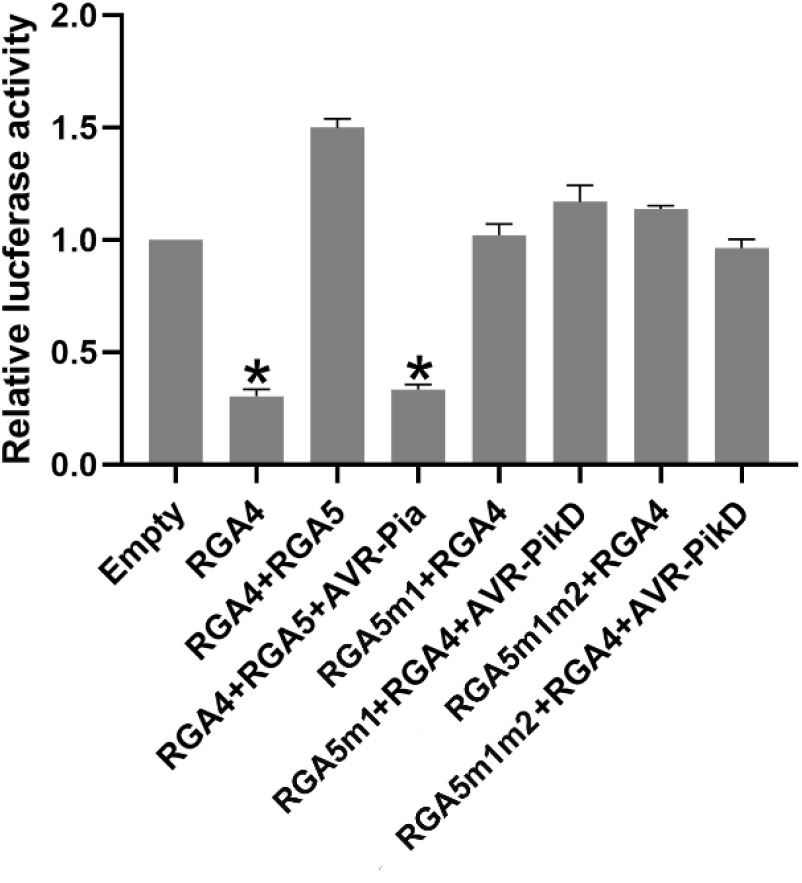
The LUC activity of rice protoplasts after transfection with different vector combinations. RGA4 and RGA4/RGA5/AVR-Pia were set as the positive control, and empty vectors served as the negative control. Significant differences with empty vector sample are labelled with an asterisk and assessed by Dunnett’s HSD test (*p*<0.05). The assays were repeated three times with similar results.

**Fig. S4.**
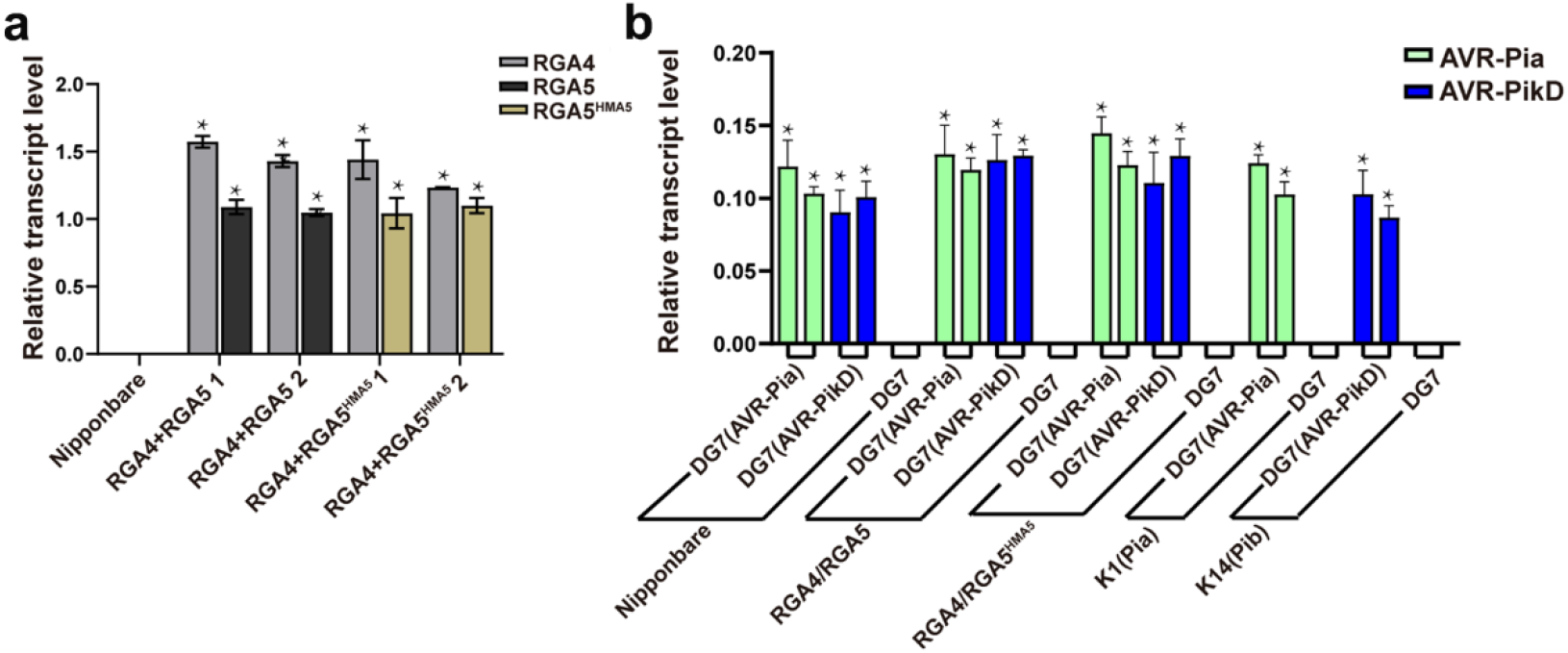
RT-qPCR analysis of the expression levels of *RGA4*/*RGA5* and *RGA4*/*RGA5^HMA5^* in two-independent transgenic cultivars (a) and *AVR-PikD* or *AVR-Pia* after inoculated onto the Nipponbare and transgenic rice leaves corresponding to the combinations in Fig.4 (b). Single asterisks represented significant differences in the expression levels (P<0.05) between wild-type and transgenic lines. The actin gene in rice or the rice blast fungus was used as the internal standard.

**Fig. S5.**
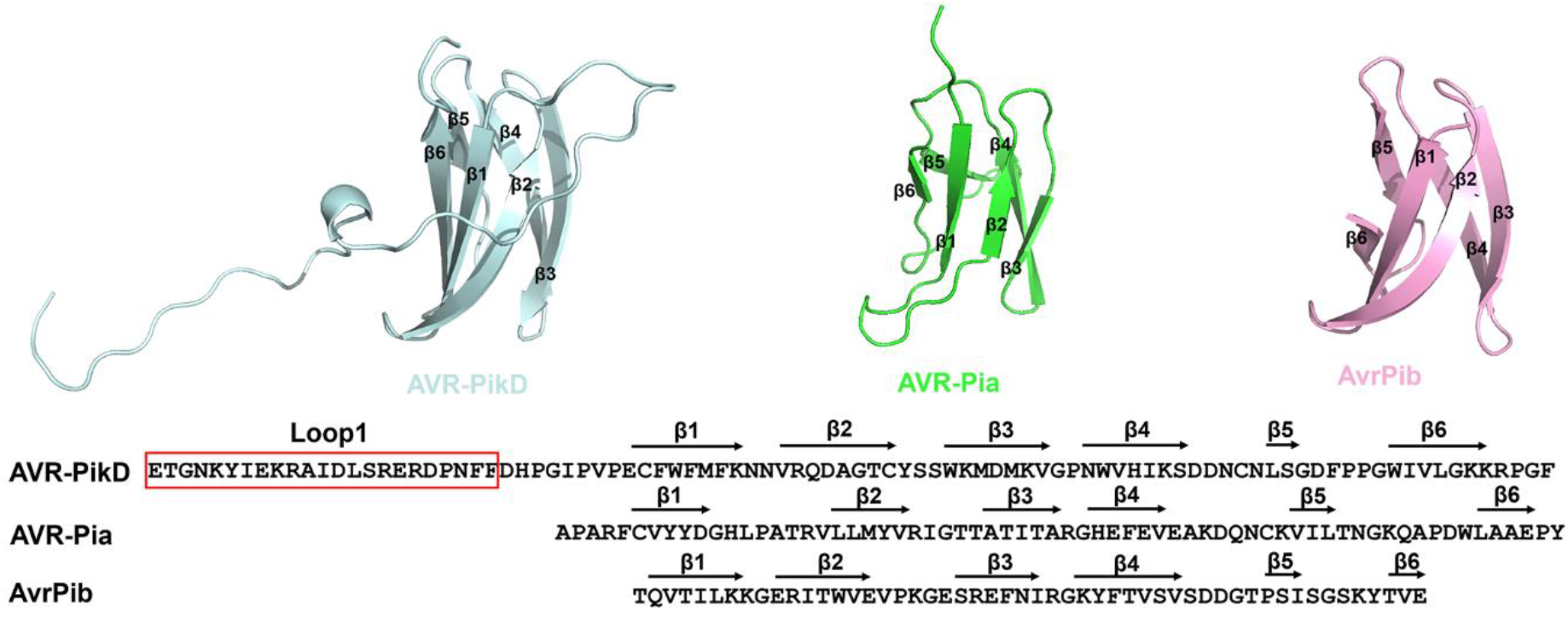
The structure and sequence alignment of AVR-PikD with AVR-Pia and AVR-Pib. Loop1 in AVR-PikD was labelled in red box.

**Fig. S6.**
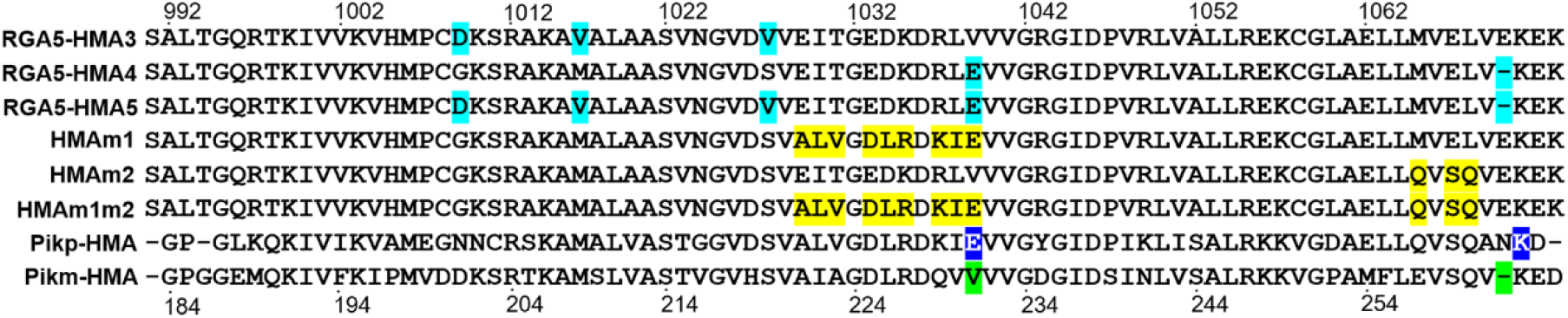
The sequence alignment of RGA5-HMA mutants with its mutants and Pikp/Pikm-HMA. The mutations in RGA5-HMA3/4/5 were highlighted in turquoise, HMAm1/HMAm2/HMAm1m2 in yellow. The key residues involved in the interaction between Piks with AVR-Pik were labelled in blue and green.

**Table S1.**
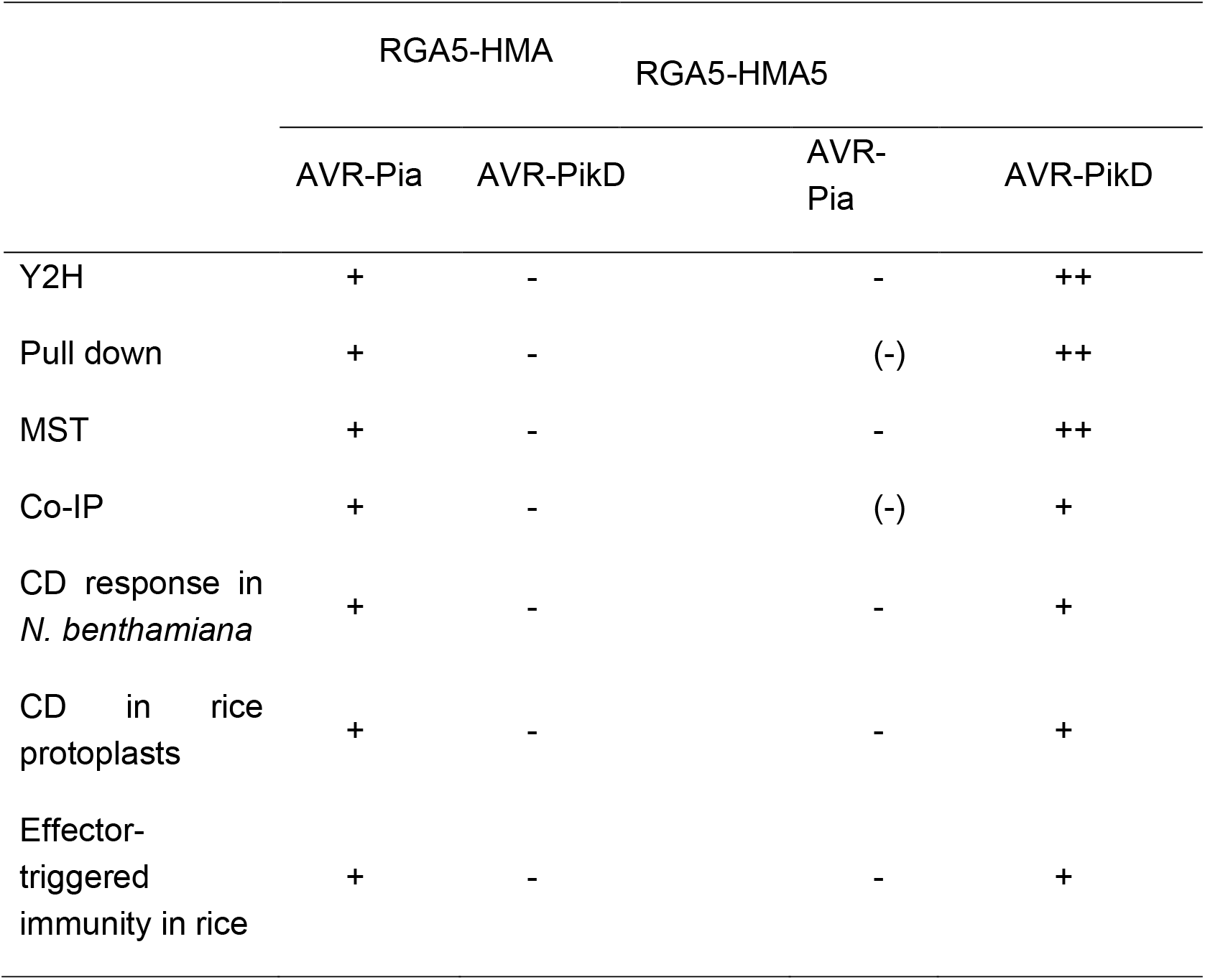
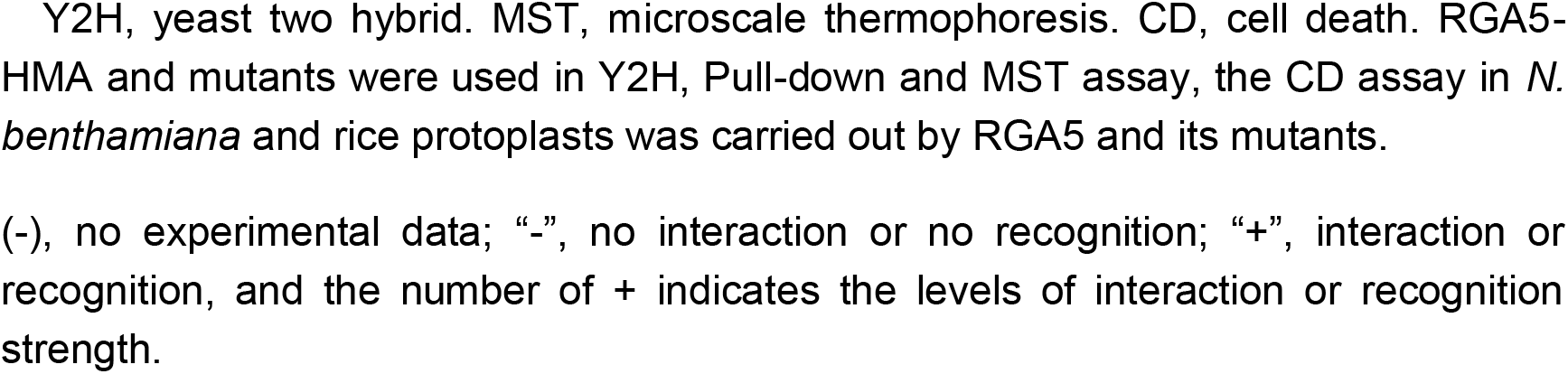
The interactions and phenotypes analysis of the HMA domain or the full-length of RGA5 and mutants with effectors

|  | RGA5-HMA |  | RGA5-HMA5 |  |
| --- | --- | --- | --- | --- |
|  | AVR-Pia | AVR-PikD | AVR-Pia | AVR-PikD |
| Y2H | + | - | - | ++ |
| Pull down | + | - | (-) | ++ |
| MST | + | - | - | ++ |
| Co-IP | + | - | (-) | + |
| CD response in<br>*N. benthamiana* | + | - | - | + |
| CD in rice<br>protoplasts | + | - | - | + |
| Effector-<br>triggered<br>immunity in rice | + | - | - | + |

Y2H, yeast two hybrid. MST, microscale thermophoresis. CD, cell death. RGA5-HMA and mutants were used in Y2H, Pull-down and MST assay, the CD assay in *N. benthamiana* and rice protoplasts was carried out by RGA5 and its mutants.
(-), no experimental data; “-”, no interaction or no recognition; “+”, interaction or recognition, and the number of + indicates the levels of interaction or recognition strength.

**Table S2.**
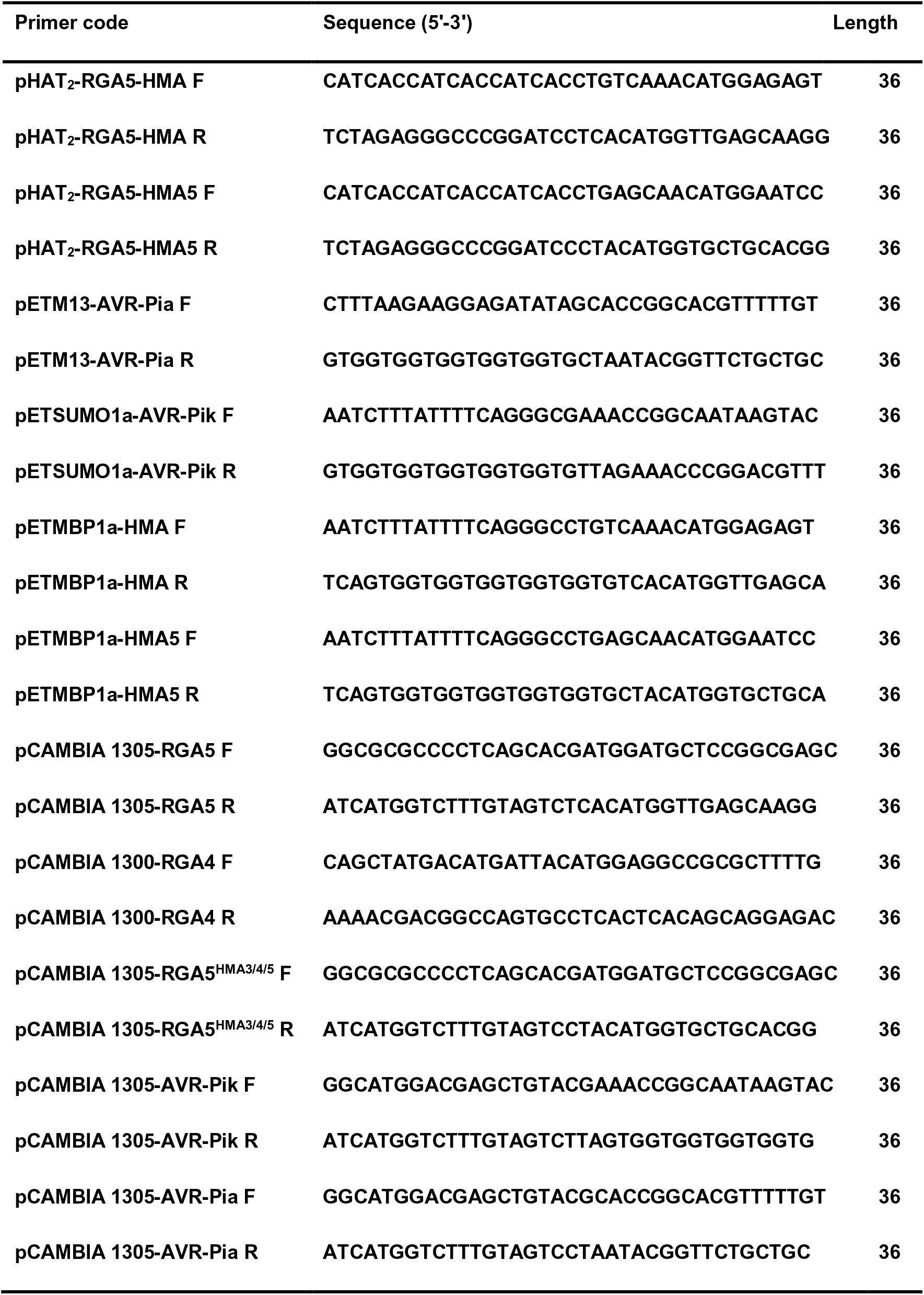
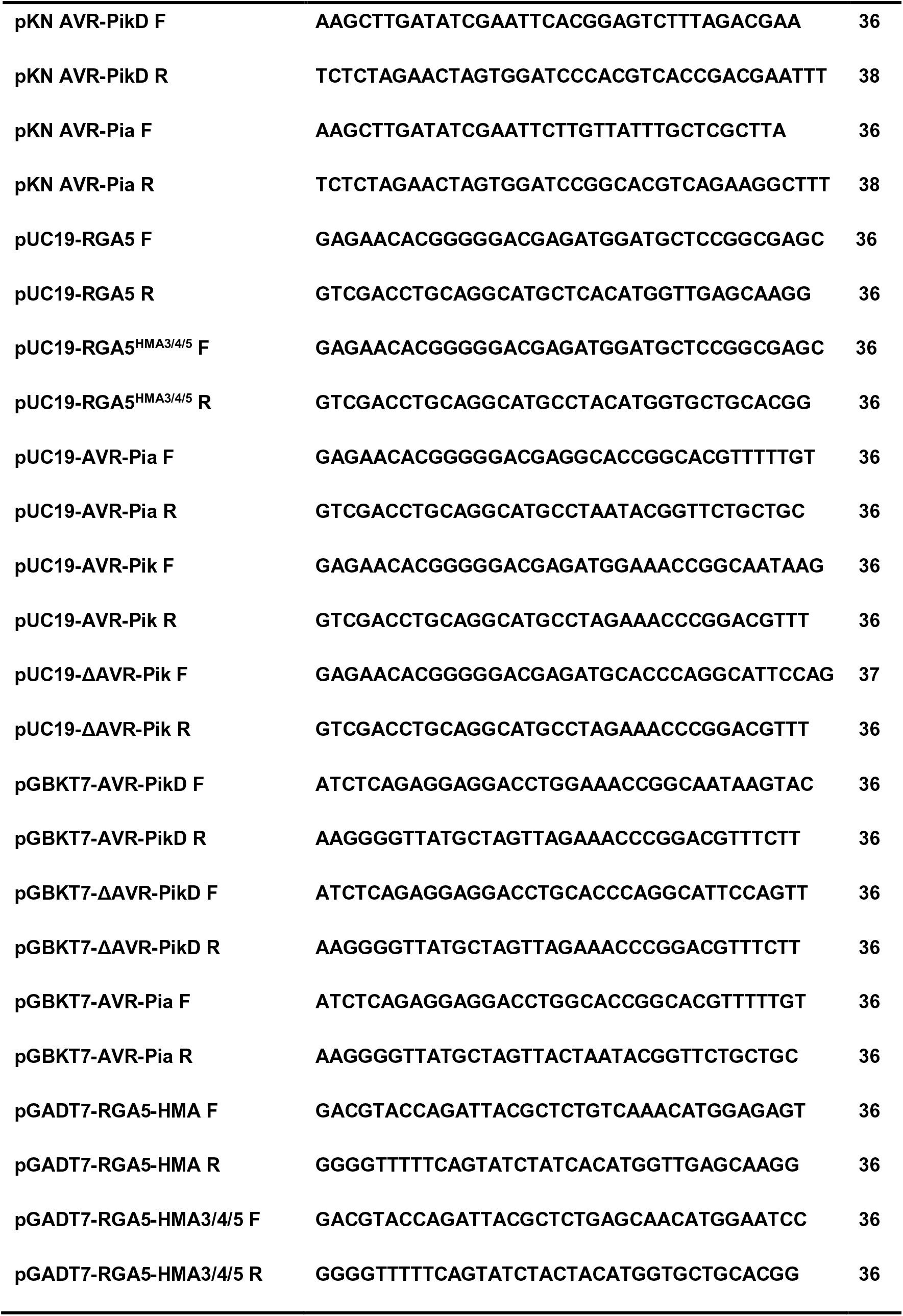
The primers used in this study.

